# Saccadic foraging reveals mechanisms of value-based decision-making in primate superior colliculus

**DOI:** 10.1101/764357

**Authors:** Beizhen Zhang, Janis Ying Ying Kan, Michael Christopher Dorris

**Affiliations:** Institute of Neuroscience, Key Laboratory of Primate Neurobiology, CAS Center for Excellence in Brain Science and Intelligence Technology, Chinese Academy of Sciences; University of Chinese Academy of Sciences, Beijing 100049, China; Department of Biomedical and Molecular Sciences, Centre for Neuroscience Studies, Queen’s University, Kingston, Ontario, Canada

## Abstract

Much progress has been made in understanding the neural basis of value-based decision-making using two-alternative forced-choice tasks. A limitation of these tasks is that the value of each option is often confounded with other cognitive factors (e.g. attention/choice probability/motor preparation). To overcome this limitation, monkeys were trained to use gaze to forage among many visual targets of varying value while neurophysiology was conducted in the saccade control center superior colliculus (SC). Surprisingly, each target’s value was represented in the SC predominantly in an absolute manner, that is, irrespective of the value of other available items. After fixating a target, the next choice was quickly selected once neural activity rose to a provisional ‘selection level’. This selection level systematically decreased, not as individual targets were harvested, but only after a class of value items was completely depleted. Our results reveal the exquisite balance across SC activities that transform value to choice.

## Introduction

Value-based decision-making involves selecting one option from many by virtue of their subjective value. Two issues are of prime importance for understanding the neural mechanism underlying such value-based decisions: 1) How is subjective value represented in the brain? 2) What is the choice mechanism that selects the one from the many? A consensus is building that regions of the frontal and parietal cortices and the basal ganglia are particularly important in calculating and representing the value of options and, perhaps, in choice selection^1–3^.

The visuo-saccadic circuit has been the primary system for studying decision-making in primates. Subjective value is computed from a number of determinants such as reward, probability and time delay in the orbitofrontal cortex (OFC)^4–6^. The OFC represents subjective value in an absolute (or ‘menu-invariant’) manner, meaning the value of each option does not depend on the other values in the choice set^7^. In addition, the OFC does not care about the spatial location of valued items. Other regions of the visuo-saccadic circuit (i.e., supplementary eye fields (SEF), lateral intraparietal area (LIP)) have clearer spatial encoding and the value of each option appears to be represented in a relative (or ‘menu-dependent) manner that depends on the value of all options under consideration^8–11^. It is unclear whether choice mechanisms ultimately act upon the absolute or relative representation of subjective value.

For the choice mechanism, two models have been proposed. One is the *goods-based model*, which holds that decisions are made in an abstract space and the choice outcome is then relayed to the premotor region which simply enacts the action for the appropriate effector^5^. The other is the *action value model*, which holds that the choice circuit encodes the subjective value associated with each potential action and a decision is made through a winner-take-all competition between these action values in the premotor circuit itself^2, 12, 13^. The question is whether these two mechanisms are exclusive or can work together to make value-based decisions.

In the current study, we focus on decision processes in the primate superior colliculus (SC). The SC is a midbrain structure that is organized as a map of potential eye movements^14^. The SC receives sensory and cognitive inputs from a large swath of the brain including the decision-related regions outlined above^15–17^. The SC exhibits sustained activity that has been linked to a number of cognitive processes, including decision making, target selection, attention, and motor preparation^18–23^. Importantly, this sustained activity may represent the last possible stage in the visuo-saccadic decision process because once the saccade threshold is reached in the SC a command is issued to the all-or-none brainstem saccade generator^24, 25^. Therefore, our goal was to understand value and choice mechanisms in this foundational brain region because it represents both the culmination of previous decision stages before it and is possibly the final arbiter of choice.

## Results

Previous research into the neural basis of decision-making has relied primarily on two-alternative forced-choice (2AFC) tasks. These tasks, however, are confounded by the fact that as the value of an option increases, the allocation of other cognitive processes associated with that option also increase (e.g., choice probability, attention, motor preparation). To overcome this confound and to examine neuronal mechanisms underlying sequential value-based decisions, we developed a novel saccade foraging task adopted from optimal foraging theory (OFT)^26^.

Two monkeys were trained to perform a saccade foraging task (Fig. 1). Anecdotally, there appears to be something particularly naturalistic and/or intuitive about this task as monkeys learn to efficiently forage in a single training session unlike many decision tasks that can take months or even years to learn. On each trial, they were presented with a rectangular array composed of visual targets of 3 equiluminant colors. Monkeys harvested reward by fixating on a visual target for a pre-specified period of time. During each block, the fixation times required to harvest the water reward associated with each color were preselected from Supplementary Table 1. Consequently, all targets of the same color were associated with a particular target value [*value = reward magnitude / fixation time*]. Monkeys were free to fixate the targets in any order they chose. Once a target was harvested of its reward, it turned into an equiluminant grey color (**Supplementary Video 1**). Between trials, the stimulus array disappeared for 3 seconds. When the array reappeared, the locations of the colored targets were shuffled but their value-color association remained intact.

**Figure 1.**
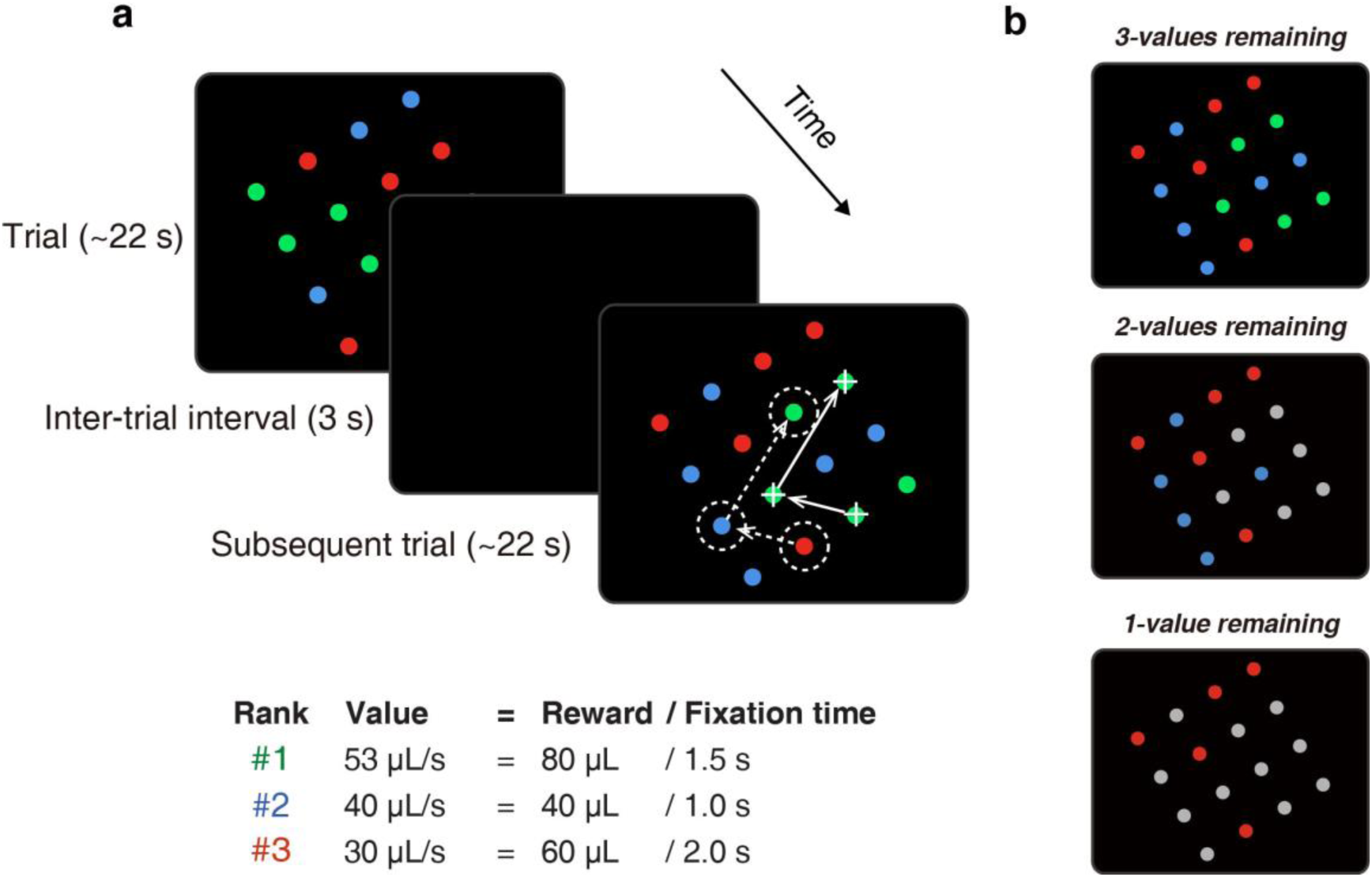
Saccade foraging task. (a) Two example trials of the saccade foraging task. The 4 × 4 array was composed of targets of 3 colors. Each target color was associated with a particular value determined by its water reward magnitude divided by the fixation time required to harvest this reward. When monkeys fixated a target for the pre-specified time, the color would turn into an equiluminant grey and corresponding reward was delivered, cueing the move to another target. In this example block, the rank of target values descended from green to blue to red. In successive trials, the association between color and value remained constant but the location of the colored targets within the array was randomized. The array size and orientation was tailored such that when the monkey was fixating a target (white cross), an adjacent target was positioned in the center of the pre-mapped response field (RF) of the isolated SC neuron (white dashed circle). The white arrows illustrate how the fovea and RF move in tandem as monkeys forage targets in the array (More detail in **Supplementary Video 1**). In a small number of experiments, larger response field eccentricities necessitated smaller 3 × 4 or 3 × 3 target arrays to fit on the visual display. (b) Examples of different menus. As monkeys tended to harvest targets in descending order of their value, the menu of items went from 3-values remaining (top), to values remaining (middle) and finally 1-value remaining (bottom).

### Monkeys were efficient value-based foragers

A critical aspect of the saccade foraging task was the monkeys were choosing under time pressure; that is, the target array disappeared before all targets could be harvested. On average, subjects were able to harvest 79 ± 6% targets. According to OFT^26^, when faced with such abundant prey items and time pressure, foragers should preferentially choose the highest valued available targets. Indeed, monkeys tended to choose scan-paths through the array in order of descending value (Fig. 2a; **Supplementary Video 1**). At the beginning of a block, monkeys often did not choose according to the OFT because they had not yet learned the association between target color and value (Fig. 2b; before dashed line). Learning tended to occur gradually over trials. But once the association between value and target color was established (i.e., ‘time of behavioral acquisition’ - Fig. 2b; after dashed line), it tended to remain stable for the remainder of the block. This pattern of behavior was not simply due to an idiosyncratic preference for a particular color because choice preference changed as the association between color and value varied between experimental sessions or, occasionally without warning, within an experimental session (Fig. 2c).

**Figure 2.**
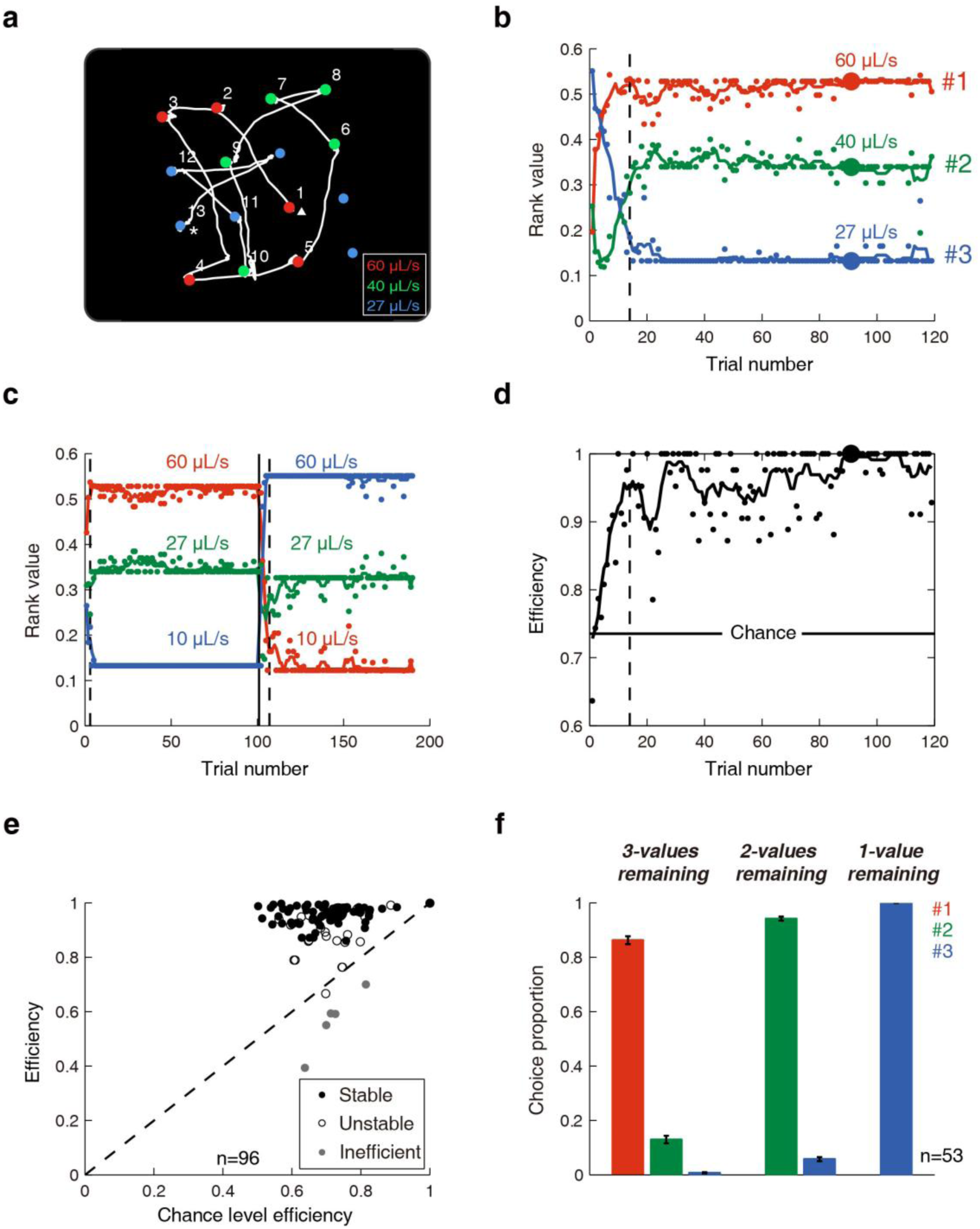
Monkeys were efficient foragers choosing targets in descending order of their value. (a) Scan path of the 91^st^ trial in a representative experiment. The white line represents the eye trace and the numbers indicate the order of successfully harvested targets. The start and end of the trial is denoted with triangle and asterisk, respectively. In some instances, such as the fixation between eye position 12 and 13, the monkey did not hold fixation long enough to successfully harvest the target. These instances were not included in subsequent analyses. The colored numbers in the legend correspond with the value of the associated target colors. This particular trial/experiment is denoted by larger data points in subsequent panels. (b) Calculating the rank value of target colors. Each dot represents the value based on the order in which a particular class of colored targets was selected within a given trial. The colored lines represent the sliding average of rank value over five trials. The dashed line represents the time of behavioural acquisition (see **Methods**) when a stable value ranking was established as determined by the Kolmogorov-Smirnov test (P < 0.05). The right colored numbers indicate the value ranking of each color which is measured from the order of average rank value across the block. (c) Same format as panel **b** except an unsignaled change in the target color-value relationship occurred at the solid line. (d) The monkey’s efficiency at harvesting water for the representative experiment shown in **b**. The black line represents the sliding average of efficiency over five trials. The horizontal line represents 95% confidence interval of chance efficiency by simulating random selection for 5000 trials. (e) Foraging efficiencies across all blocks plotted against their corresponding chance efficiencies. Only experiments that displayed significantly efficient and stable preference (black filled dots) were included in further analyses whereas inefficient (gray filled dots) and unstable blocks (unfilled black dots) were excluded. (f) Choice preferences as menu transitioned from 3-values, to 2-values, and to 1-value targets remaining. Error bars represent ±SEM. (***N.B.*** - The association between target color and value ranking was randomized between each experiment. However, for display purposes, red, blue and green will indicate the number 1, 2 and 3 value rankings, respectively, throughout the remainder of the paper.)

As monkeys learned to choose targets in descending order of value, their efficiency at harvesting water also increased (Fig. 2d). We defined the efficiency as the value of each chosen target divided by the highest value that was available within the array at that time. Across experimental sessions, monkeys’ efficiency after behavioral acquisition was better than chance (Fig. 2e; paired-t test, P = 7.3 × 10^-27^). Once behavioral acquisition was reached, the value rankings tended to remain stable thereafter (Fig. 2e - filled data points). The sessions with unstable value preferences (Fig.2e - unfilled dots, see Methods) and the five sessions during which efficiency was significantly lower than chance (Fig. 2e - grey dots) were excluded from further analyses. Average efficiency of the remaining sessions was 0.96 ± 0.03. All subsequent analyses were performed only on those trials after time to behavioral acquisition was reached (see **Methods**).

According to OFT, monkeys should harvest prey items based on the *relative* value of all available prey items; therefore, once a certain class of prey items is fully harvested, preferences should change accordingly. Indeed, monkeys preferentially chose the highest value targets for a given menu (Fig. 2f). After the most valuable targets were exhausted (Fig. 2f; red), monkeys chose the second most valuable targets (Fig. 2f; green), and after those were exhausted, they chose the least valuable targets (Fig. 2f; blue). We hypothesize that there would be a marked change in SC representation of target items associated with these menu changes.

### The SC encodes both value and choice

We recorded 96 single neurons in the intermediate and deeper layers of the SC from two monkeys during the saccade foraging task. In total, 53 neurons satisfied our neurophysiological criteria while the monkeys’ performance simultaneously satisfied our behavioral criteria to be included in the following analyses (see **Methods** and Fig. 2e).

SC activities were correlated with value ranking of targets in the neuronal response fields (RF; Fig. 3a). To be clear, the order in which the foveae harvested targets provided us with a behavioral measure of the subjective value ranking of the colored targets (Fig.2), however, each recorded SC neuron analyzed a target *adjacent* to the current foveae location and in its RF (see gray shading in **Supplementary Video 1**). Whereas the foveae harvested targets in a fairly strict descending order of their value, the targets that entered the neuron’s RF was fairly random. Within each menu, the neuronal activity increased as the value of the target in the RF increased (Fig. 3a). Moreover, if a previously harvested grey target entered the RF, this elicited the least SC activity. The latter may represent the baseline sensory activation because these harvested targets had no value and were virtually never the target for a saccade. However, neuronal activity was not only modulated by value in the RF, but was also highly predictive of choice. Neuronal activity leading up to choices directed towards the RF target (Fig. 3b; Choice-in) were significantly stronger than that preceding a choice to targets outside the RF (Fig. 3b; Choice-out).

**Figure 3.**
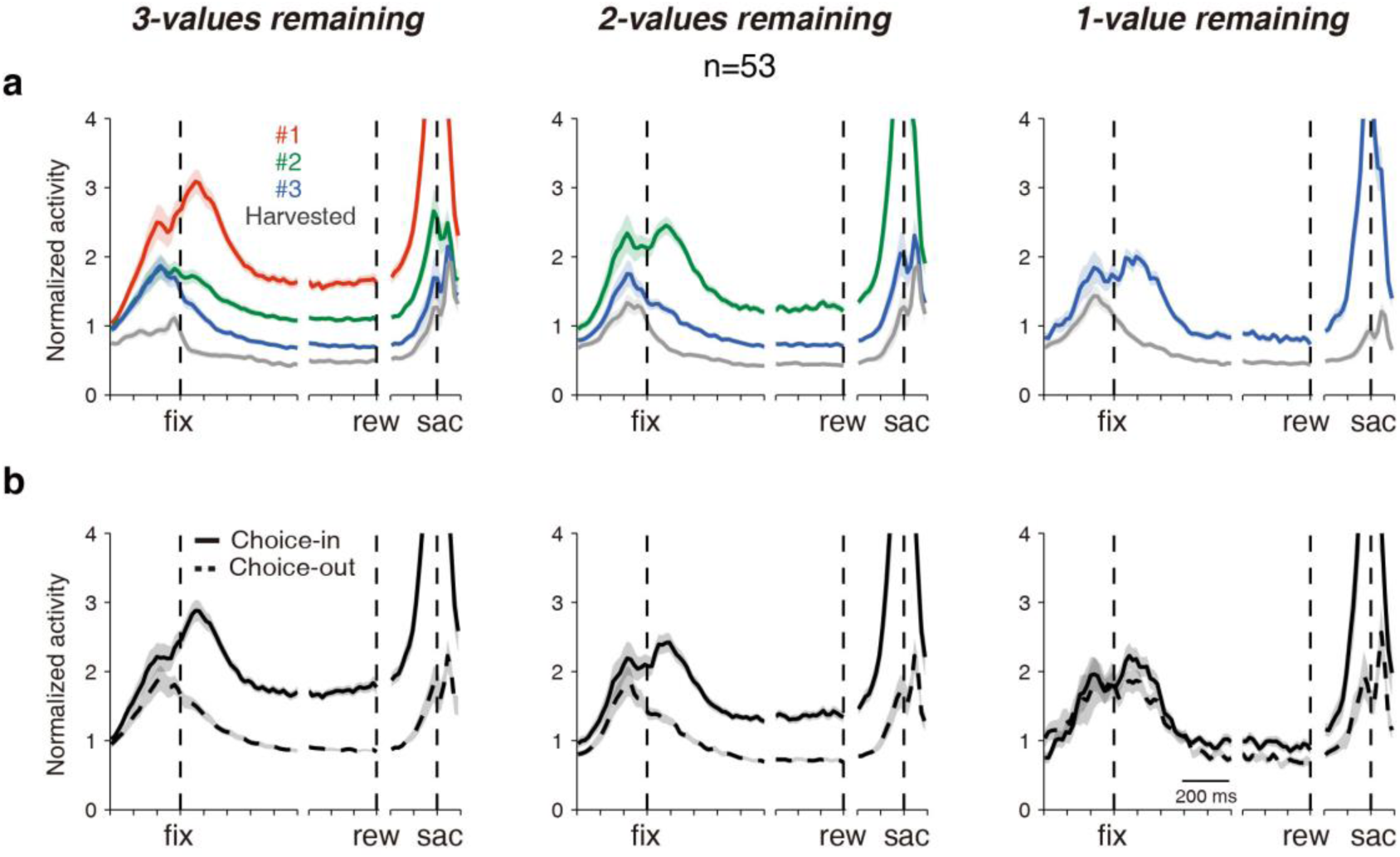
Population neuronal activity reflected target value in the response field and predicted upcoming saccade choice. Note that plots represent (a) the value of, or (b) choices directed into or out of the response field target rather than properties of the currently fixated target. The left side of each panel is aligned on the beginning of fixation (fix), the middle is aligned on reward delivery (rew), and the right is aligned on saccade onset (sac). The total duration of the spike density waveforms could not be shown because the fixation time varied across targets. The shaded regions surrounding each line represent SEM.

This analysis – similar to those used in most 2AFC tasks – is difficult to interpret because monkeys are more likely to choose high valued targets (see Fig. 2f); thus, value and choice encoding cannot be disentangled. A major advantage of our saccade foraging task is that there are many potential targets that share the same value but are not necessarily the target of next choice which allows us to dissociate the contribution of each to neuronal activity.

After the variables were isolated from each other, SC activity still remained correlated with both target value in, and upcoming choice towards, the RF (Fig. 4a); although each displayed a different time course. Multiple linear regression analyses showed the value signal began to significantly increase approximately 300ms before the target was even fixated (i.e., Fig. 4b; horizontal black lines) and peaked within 200 ms of a new target entering the response field. Perhaps this pre-fixation activity represented predictive ‘remapping’ of value signals akin to the sensory remapping that occurs when saccades will bring visual stimuli into SC response fields^27, 28^. In contrast, the choice signal increased significantly only after the fixation period began (Fig. 4b; horizontal gray lines). Overall, there was a ‘value-to-choice’ transformation such that value signals were dominant early during a new fixation and choice signals became dominant as the fixation period progressed (Fig. 4b). We will describe this transformation in detail below.

**Figure 4.**
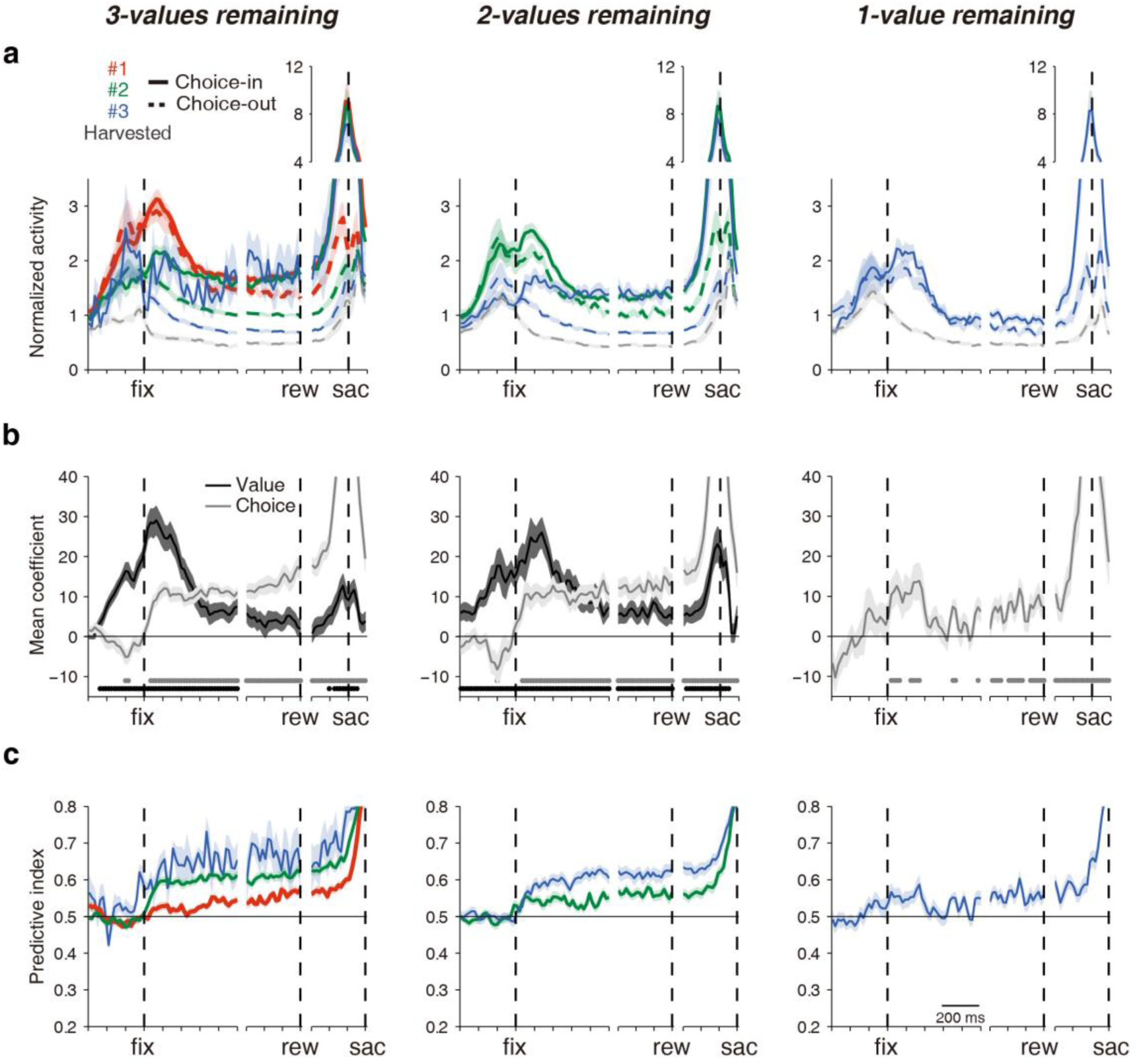
The evolution of value and choice signals in SC activity across 3 menus. (a) Normalized neuronal activity was segregated based on the value of the target in the response field (line color) and whether a saccade was ultimately directed to the response field target (solid lines) or a target outside the response field (dashed lines). Otherwise, the same format as Fig. 3. (b) The evolution of the population regression coefficients for value ranking (black) and choice direction (gray). There is no value plot in the rightmost 1-value panel because all remaining targets had the same, lowest value. The time point with significant regression coefficients by value ranking (black) and choice direction (gray) was shown with dots at the bottom of the panel (t test, P < 0.05, with False Discovery Rate correction). (c) The evolution of neuronal choice selectivity towards targets of different value rankings. The receiver operating characteristic analyses were done between activities associated with choices toward the response field target versus targets outside the response field that shared the same value. All shaded regions represent SEM.

### The Evolution of SC Value and Choice Signals

Slightly before and during fixation of a new target, neuronal activity primarily reflected the value ranking of the target in its RF. Immediately upon establishing fixation, however, the choice signal quickly began to ramp up (Fig. 4b). As the fixation period progressed, there was a marked differentiation in SC activity based on whether the monkey would ultimately look to the RF target or elsewhere.

A stable, common level of activity was reached in the SC if the monkey would ultimately choose the RF target regardless of its value (Fig. 4a; clearer in Fig. 5a). Within each given menu, the choice-in neuronal activity of each value ranking reached a common level during the late fixation period (Fig.5b; n-way ANOVA test, effect of value, F_2, 250_ = 0.31, P = 0.73). We named this common choice-in level of activity the ‘*selection level*’ that, as will be evidenced below, represents provisional choice. Consistent with this proposal, both multiple linear regression (Fig. 4b) and signal detection (Fig. 4c) analyses showed that choice signals build up and reach a plateau shortly after a new target is fixated. Moreover, behavioral results show that, within a given menu, choice-in saccade latencies were the same regardless of the value of the saccade targets (Fig. 5c; n-way ANOVA test, effect of value, F_2,258_ = 0.01, P = 0.99). This is consistent with a neural signal that starts for a common level ramping up to a saccade bound.

**Figure 5.**
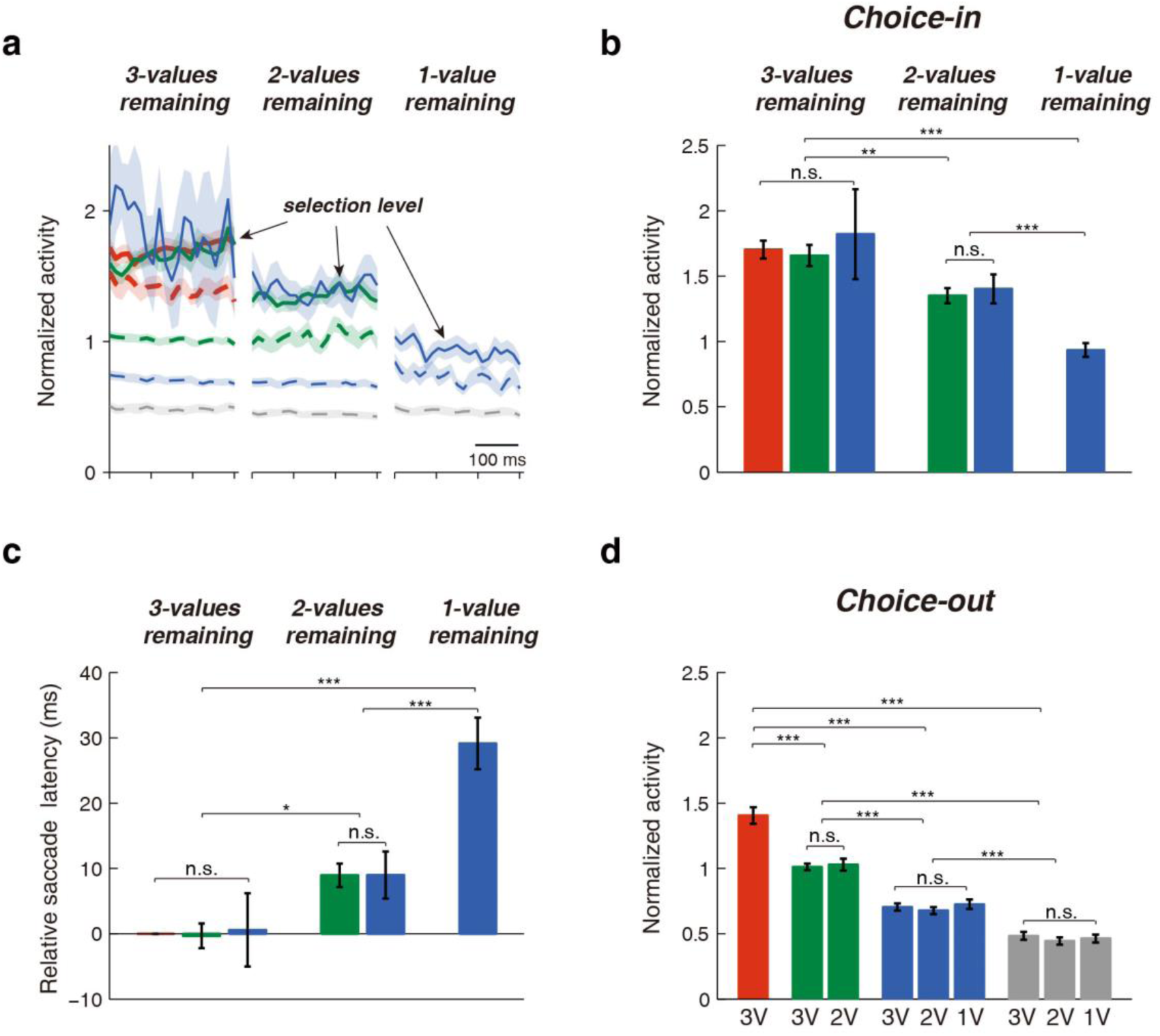
Menu updating of the selection level and value representations. (a) Normalized neuronal activity during the last 300ms of fixation. Same format as Fig. 4a. (b) Mean activity ± SEM associated with choice-in conditions. (c) Relative saccade latencies towards different value ranking targets across the 3 menus. All latencies were relative to the 1^st^ value ranking targets in the 3-values remaining menu. (d) Mean activity ± SEM associated with choice-out conditions. Data is presented for each value ranking when there were three values (3V), two values (2V) or one value (1V) targets remaining in the array. (n.s., nonsignificant,*P < 0.05, **P < 0.01, ***P < 0.001; n-way ANOVA tests, *post hoc* tests were done with Bonferroni correction)

Conversely, if the RF target was not selected for the subsequent saccade, SC activity remained strongly correlated with its value (Fig. 5a; Fig. 5d; n-way ANOVA test, effect of value, F_3, 449_ = 174.93, P = 4.1 × 10^-75^; note all the significant comparisons between values shown in Fig. 5d).

### Menu updating of the selection level

To understand how decision circuits accommodate menu changes, we compared SC activity across menus (Fig. 5a). Although the selection level remained constant within a menu regardless of the value of the chosen target, it systematically decreased across menus (Fig. 5b; n-way ANOVA test, effect of menu, F_2, 250_ = 16.86, P = 1.4 × 10^-7^). This decreasing selection level was not the result of harvesting individual targets (Supplementary Fig. 1a; paired-t test; 3-values, P = 0.083; 2-values, P = 0.72; 1-value, P = 0.067) but occurred during the transition between menus after entire value classes were harvested. Nor can it be explained by changes of the value of fixated targets rather than RF targets (Supplementary Fig. 1b; 3-values, one-way ANOVA, F_2, 99_ = 0.85, P =0.43; 2-values, paired-t test, P = 0.097). These menu-dependent changes in the selection level were reflected in the saccade latency (Fig. 5c; n-way ANOVA test, effect of menu, F_2, 258_ = 23.8, P = 3.3 × 10^-10^). As the selection level decreased with fewer menu items, it presumably got further from the hard saccade threshold in the SC^14, 29^ resulting in saccade latency that increased as the menu items decreased from three to two to one. Together, this suggests that the activity level that constitutes preliminary choice selection is dynamically updated across the SC map with changes in menu items.

### Menu updating of value coding

Across menus, the value signal differed during the early versus late fixation period. During late fixation, the choice-out activity of each value ranking remained unchanged across menus (Fig. 5d; n-way ANOVA test, effect of menu, F_2,449_ = 0.27, P = 0.76). In contrast, the initial visual response after fixating a new item was menu-variant with respect to value (Fig. 4a; clearer in Supplementary Fig. 2). During early fixation, the choice-in activity of valued targets (Supplementary Fig. 2a; 2nd rank, 3-values versus 2-values, paired-t test, P = 0.0012) and choice-out activity of valued targets (Supplementary Fig. 2b; 2^nd^ rank, 3-values versus 2-values, paired-t test, P = 4.0 × 10^-9^; 3^rd^ rank, one-way ANOVA tests, F_2,142_ = 18.29, P = 8.5 × 10^-8^, *post hoc* tests, 1-value versus 3-values, P = 7.8 × 10^-7^, or 2-values, P = 1.1 × 10^-6^) scaled according to the other options available in the menu.

### Balanced activity between selected and unselected options

An upshot of having a menu-variant selection level but menu-invariant value representations is that choice selectivity remained remarkably consistent under all conditions. When we performed signal detection analyses for the selected target versus the highest available non-selected representation on the SC map, the neuronal selectivity remained approximately 55% during the steady-state late fixation period (Fig. 6a). This 55% neuronal selectivity remained unchanged regardless of the value of the chosen target or the composition of the targets in the menu (Fig. 6b). This result suggests that there may be an inherent equilibrium achieved between the activations for the selected and non-selected targets representing provisional choice that can be maintained for extended periods on the SC map.

**Figure 6.**
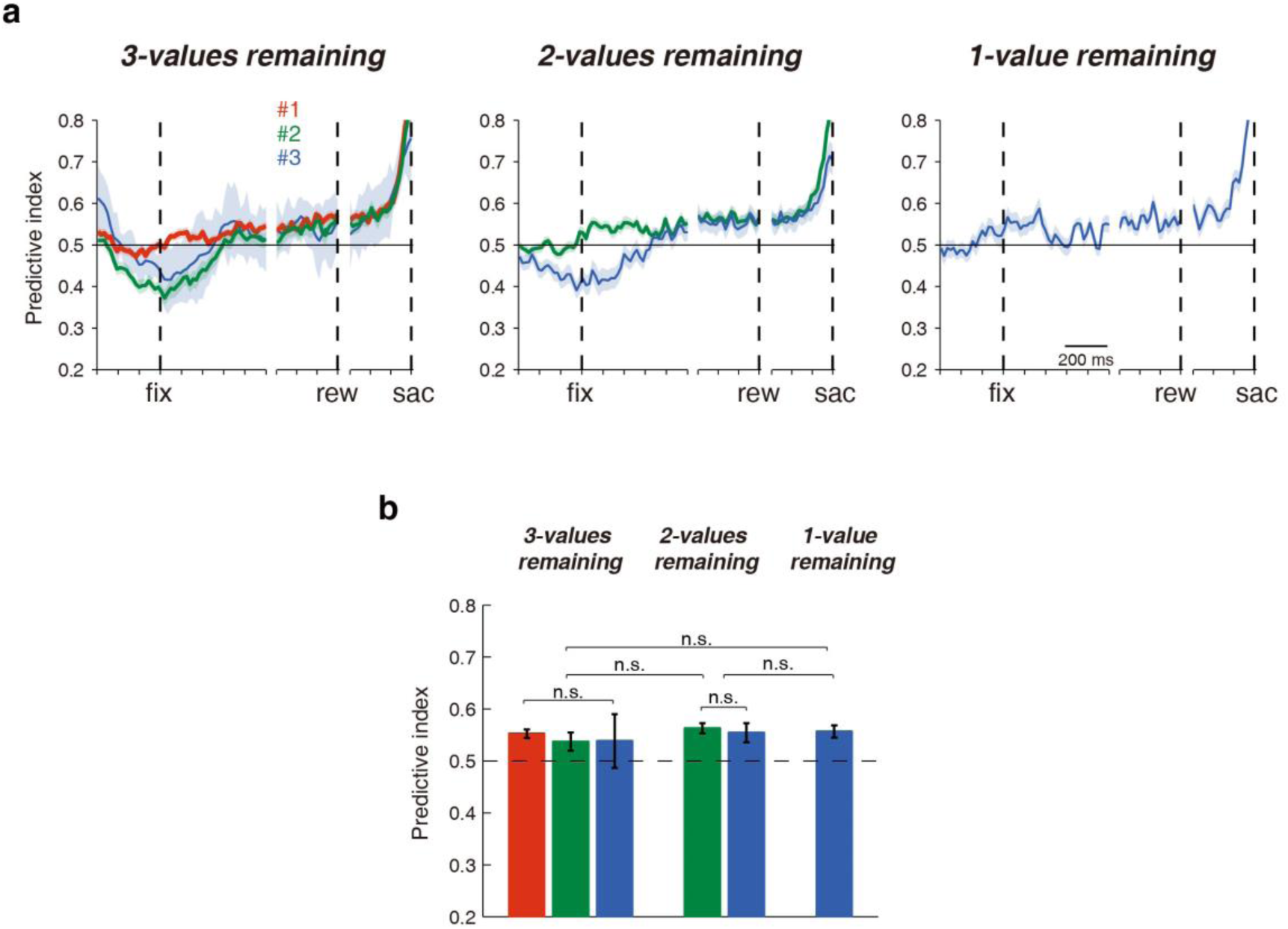
The selectivity between selection level activity and the highest available choice-out activity remained unchanged across menus. (a) The evolution of neuronal choice selectivity when comparing choice-in activities of each value ranking with the highest available choice-out activity. All shaded regions represent SEM. (b) The average predictive indexes during the late fixation period from **a**. Error bars represent ± SEM. Data is presented for each value ranking when there were three values (3V), two values (2V) or one value (1V) ranking targets remaining in the array. (One-way ANOVA tests)

### Manipulating SC activity biased choice in a manner predicted by SC recordings

Previous research suggests that action selection could happen through the competition of multiple movement plans in the premotor regions^2, 13^. As both the selection level and the value-modulated activities of unselected targets were represented simultaneously on the SC maps, we hypothesized that a competitive action selection process happened during the late fixation period. We tested this hypothesis by perturbing activity in the late fixation period with electrical micro-stimulation in an attempt to bias choice patterns. More specifically, we hypothesize that the efficacy with which micro-stimulation biases choice is a function of the neuronal distance between the value activity associated with the target at the stimulation site and the menu-dependent ‘selection level’ (i.e., the distance between the relevant dashed line and solid lines in Fig. 5a).

Our goal was to bias competing selection processes on the SC map without directly triggering a saccade itself, so we used sub-threshold electrical micro-stimulation to increase SC activity^18^. Two lines of evidence suggest that micro-stimulation was sub-threshold. First, we only included experiments if no saccades were triggered during the micro-stimulation period (Supplementary Fig. 3a; bottom panel grey box). Second, we did not observe any difference in saccade latency between stimulation and control conditions both in the example block (Supplementary Fig. 3b; Wilcoxon rank sum test, P = 0.60) and across the population of stimulation sites (Supplementary Fig. 3c; choice-in, paired-t test, P = 0.34; choice-out, paired-t test, P = 0.25).

Next we assessed whether this sub-threshold, micro-stimulation could affect choice. The proportion of choices directed towards the stimulation site increased significantly after stimulation compared to the non-stimulation control condition (paired-t test, P = 1.5 × 10^-12^; not shown). However, micro-stimulation did not bias choices towards all targets equally. Micro-stimulation biased choices predominantly when there were high value targets at the stimulation site (Fig. 7). When there were 3 values in the menu, micro-stimulation significantly biased choices towards highest valued targets (mean = 0.104 ± 0.013, paired-t test, P = 4.3 × 10^-11^), less so to middle ranking targets (mean = 0.053 ± 0.009, paired-t test, P = 1.7 × 10^-7^), and not at all to lowest valued targets (mean = 0.000 ± 0.002, paired-t test, P = 0.9). Once the highest valued targets were all harvested, the 2^nd^ ranked targets became the most valuable, and micro-stimulation exerted a stronger biasing effect towards them than when there were 3 menu items remaining (mean = 0.083 ± 0.014, paired-t test, P = 1.2 × 10^-7^). Only when there was 1-value remaining in the menu did micro-stimulation exert a significant biasing effect towards the target at the stimulation site (mean = 0.105 ±, paired-t test, P = 2.5 × 10^-4^). Therefore, the overall biasing effect of micro-stimulation on choice was both value- and menu-dependent (Fig. 7b; n-way ANOVA; effect of value, F_2, 427_ = 19.6, P = 7.2 × 10^-9^; effect of menu, F_2, 427_ = 17.2, P = 6.7 × 10^-8^) and mirrored very closely the value- and menu-dependent pattern of behavioral choice (Fig. 2f) and SC activities (Fig. 5a). Together these results suggest SC mechanisms are involved in the action selection process of value-based decision-making.

**Figure 7.**
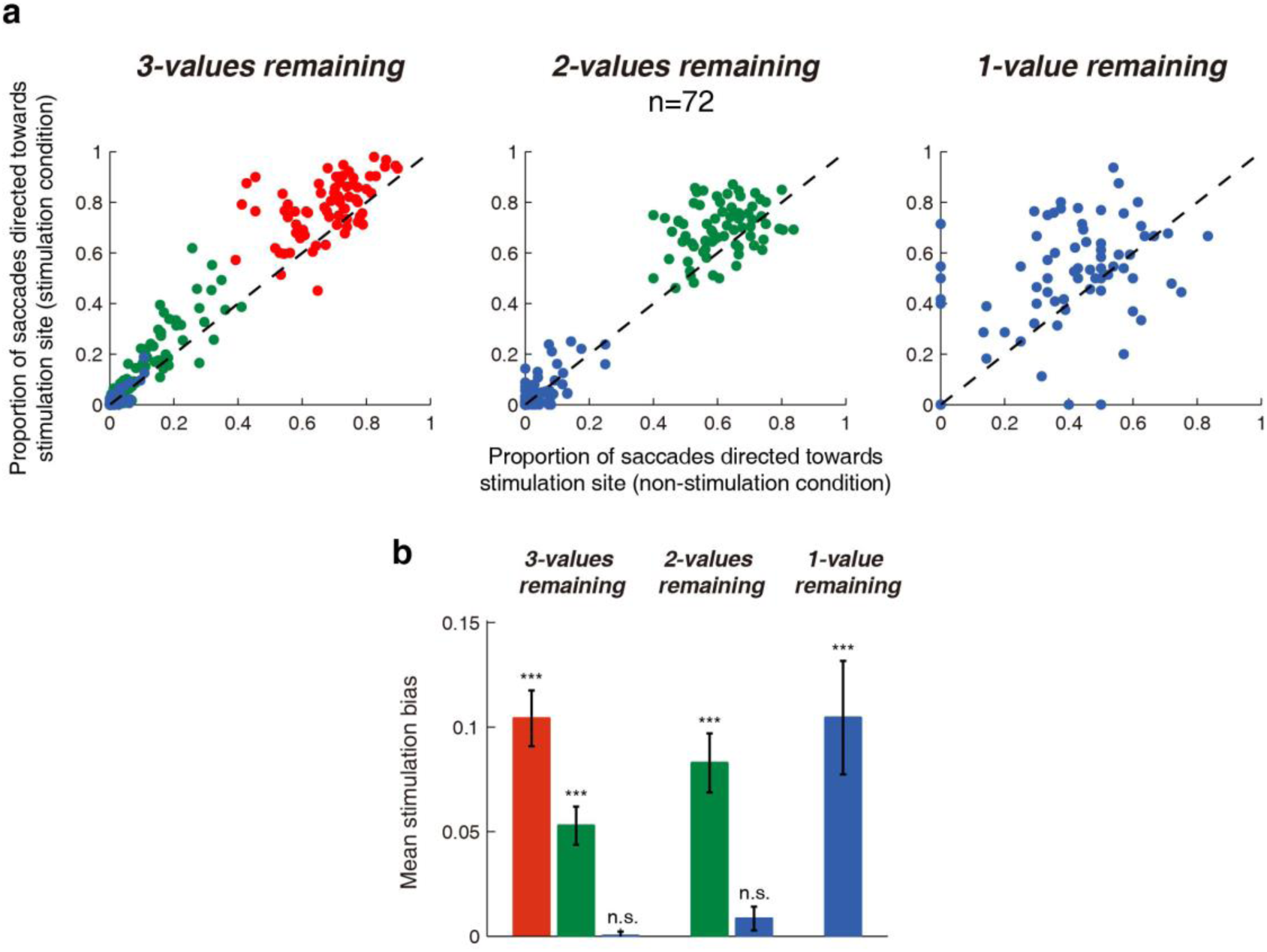
Sub-threshold micro-stimulation applied during the late fixation period biased choice. (a) The proportion of choices directed towards the stimulation site target in stimulation condition vs. non-stimulation control condition across the 3 menus. Red, green and blue denote when 1^st^, 2^nd^ and 3^rd^ value ranking targets were located at the stimulation site respectively. The dashed line represents the line of unity. (b) Difference in the proportion of saccades directed towards the stimulation site under stimulation condition minus non-stimulation condition. Error bars represent ± SEM. (Paired-t tests)

## Discussion

We employed a novel foraging task to examine the role of the primate SC in choosing value-based saccades. Multiple targets sharing the same value and the numerous successive choices afforded by this task allowed us to examine neural processing both when value was dissociated from choice and when changes occurred in the menu of value items. Monkeys performed this task at a near optimal level as predicted by optimal foraging theory (Fig. 2). At the neuronal level, we observed a transformation from value-to-choice in SC processing (Fig. 4) and choices were altered using sub-threshold, electrical micro-stimulation (Fig. 7). These findings suggest that the SC is an active participant in value-based saccadic decision-making. Three additional novel - and somewhat surprising - findings were observed. First, value was represented in an absolute or menu-invariant manner across the SC map interrupted only briefly by relative value representations (Fig. 8a). Second, activation at one location on the SC map was elevated to a common ‘selection level’ that predicted upcoming choice regardless of its associated value (Fig. 8b). Third, this common selection level was menu-dependent decreasing as classes of menu items were removed from the choice array (Fig. 8c). We will illustrate each of these novel findings in turn using the schematic model shown in Figure 8.

**Figure 8.**
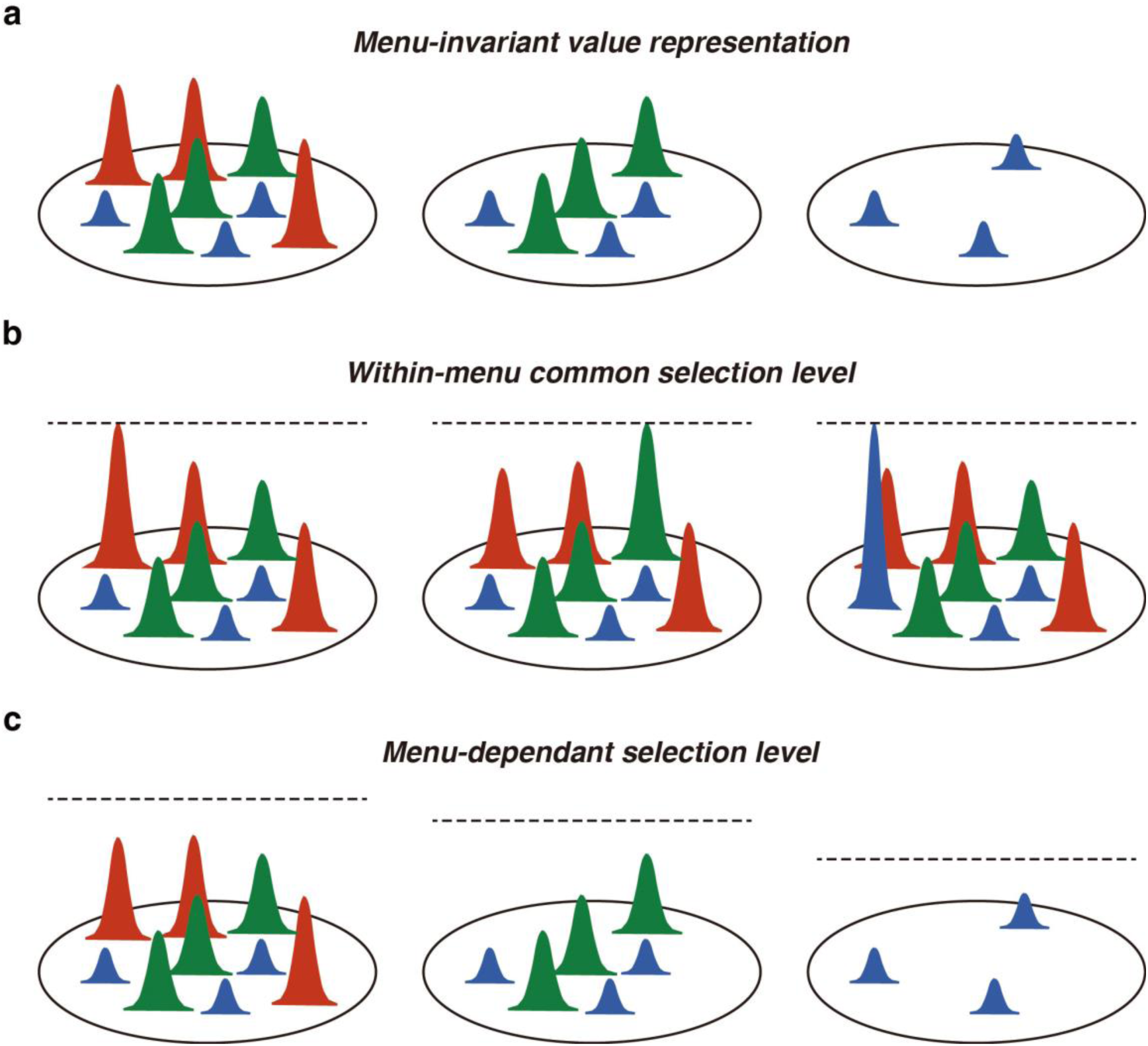
Three SC mechanisms important for foraging decisions. The Gaussian curves represent late fixation population activities on the SC map associated with the highest (red), middle (green) and lowest (blue) valued targets in the visual array. The dashed line represents the selection level. (a) The value of targets was represented across the SC in an absolute manner that did not vary as the menu decreased from values remaining (left) to 2-values remaining (middle) to 1-value remaining (right). (b) Within the same menu, neuronal activity associated with the selected option reached the same “selection level” (dashed line) regardless of whether the highest (left), middle (middle) or lowest (right) valued target was chosen. (c) The selection level decreased as the menu changed from 3-values remaining (left), 2-values remaining (middle) and 1-value remaining (right).

### Both absolute and relative value representations in the SC

There has always been a mystery surrounding how value is represented in the brain at the most fundamental level. On one hand, convincing evidence suggests that value is represented in a menu-invariant or absolute manner in the OFC albeit in a highly non-spatialized manner^7^. On the other hand, equally convincing evidence suggests that value is represented in a menu-dependent or relative manner in more spatially organized visuo-saccadic regions like LIP^8–10^. Here we demonstrate that as you approach the end of the saccadic decision pathway in the SC, value dynamically alternates between absolute and relative representations.

A menu-invariant, absolute representation of value dominated the SC map in our experiment. In fact, except for a few hundred milliseconds surrounding each new fixation (see below), it was absolute value that was represented through the duration of a multiple-hour experiment. Unlike absolute value representations observed previously in OFC, however, this SC representation was spatially organized across a topographic map in a manner more appropriate for driving motor action. Figure 8a shows the late fixation period activation associated with items of different value. This activation was menu-invariant because it remained the same for each value item regardless of the number of menu items available (Fig. 5). Ultimately, decision-making models require a comparison between absolute values because we often need to compare items across menus and absolute value ensure preference is transitive (i.e., if one prefers A >B, and B>C, this implies A>C)^5, 30, 31^, which is a hallmark of rational choice behavior^32^.

However, like other visuo-saccadic brain regions, we also observed relative value signals in the SC but only during the initial visual transient surrounding a new target landing in the RF (Supplementary Fig. 2). It has been proposed that relative value signals may function to normalize neural activity to rescale absolute value such that differences between available options are maximized leading up to choice^2^. Indeed, it appeared that it was during this transient relative value period that the decision was made because this is when value was transformed into choice and subsequently maintained at a relatively steady-state selection level (Fig. 4). The menu-modulated visual transient may also perform a function related to attention or saliency associated with the target that has recently entered the neuron’s response field ^33–35^.

### The selection level represents provisional choice

Our second major finding is that activity associated with the target for the next saccade reached a heightened selection level (Fig. 8b). Any selected target reached the same level of activation regardless of the value of the item. Furthermore, this selection level was maintained at a steady state until the cue for a movement acted to trigger the subsequent motor burst. Although one target was selected with heightened activity, the non-selected options maintained their absolute value representations across the SC map.

We believe that the selection level represents a provisional decision that was held until the proper time of motor execution arrived. We use the term “provisional” because when artificial stimulation was applied to the SC after the selection level was established, choice probability could still be altered (Fig. 7). Therefore, our results suggest the selection level represents the current provisional decision but the ultimate action does not occur until the putative ‘hard’ bound is crossed in the SC^14, 29^ (e.g., Fig. 4 – ‘sac’ period). In our task, the relationship between target color and value was fixed throughout. If, however, value information could change during the fixation period, the target for the next saccade presumably could still be altered. Similarly, a relatively common level of activation has been observed in perceptual decision tasks when a selected option must be held for a fixed duration before execution regardless of the strength of evidence for that option^22, 36^. This suggests that the selection level may represent provisional choice across a wide range of decision tasks.

### The selection level changes across menus

Our third finding is that the selection level systematically decreased with fewer menu items (Fig. 8c; see decreasing dashed line across menus). This was not simply a matter of fewer available targets but occurred abruptly once an entire class of value items was fully harvested (Supplementary Fig. 1a). The upshot of this dynamic selection level was that the “distance” between the selection level and the presumably fixed saccade threshold also varied across menus. This variability in neural distance across menus accurately predicted subsequent saccadic reaction times (Fig. 5c) and the ability of micro-stimulation to bias choice towards the stimulation site target (Fig. 7). Moreover, the distance between the selection level and the next highest available item on the SC map remained remarkably consistent across conditions as evidenced by constant neuronal selectivity (Fig. 6).

At this point it is not clear why or how the selection level should decrease in this manner nor how the exquisite balance between selected and non-selected representations is maintained. Efforts to model the competitive interactions across the SC map when faced with a large array of valued items may shed some light on this issue^37, 38^. But the fact that the selection level is adjustable under the current conditions implies that the selection level may also be adjustable under other conditions such as speed-accuracy tradeoffs^39, 40^ or confidence estimations^41, 42^.

### Evidence for action value model in the SC

Once the selection level has been established, the pattern of activities across the SC closely follows that proposed by action value models^2^. Action value models propose that decisions are made through the competition of movement plans. Indeed, our results satisfy the criteria outlined by Padoa-Schioppa for an action value signal to be involved in economic decisions^5^. First, activity must be motor in nature. Second, activity must be modulated by subjective value. Third, activity must not be downstream of the decision. The SC is undoubtedly motor in nature^14^, we demonstrate that SC activity is strongly modulated by value (Fig. 4) and, to our knowledge, our demonstration that choice can be biased using sub-threshold micro-stimulation (Fig. 7) is the first to satisfy the third criteria. The spatial organization and competitive interactions across the SC map make it ideally suited to enact a winner-take-all mechanism to rapidly take the provisionally selected target to the final motor action once the movement cue is given^43, 44^.

Given the ample evidence to show that a goods-based value mechanism is employed in the OFC, we surmise that this provides the inputs for the absolute value signals observed in the SC topographic map. However, given the topographic mapping, close relationship with the motor plant, competitive interactions between population activities across the SC, and causal stimulation evidence presented above, we propose that it is an action value mechanism that selects the one from the many and executes the final choice. Of course, here subjects are making saccadic movements to targets located in a stable array. Under conditions in which the spatial location of valued items or the effector needed to acquire them is less explicit^45^ – that is, more abstract decisions – presumably a goods-based neural mechanism would dominate.

## Methods

### Animal preparation

Two male rhesus monkeys (Macaca mulatta; D, 12 kg, and R, 8 kg) participated in this study. Physiological recording and stimulation techniques as well as surgical procedures were described previously^18, 46^. Animals were under the close supervision of the Institute veterinarian. All procedures were approved by the Animal Care Committee of Shanghai Institutes for Biological Sciences, Chinese Academy of Sciences (Shanghai, China).

Throughout the experiments, monkeys were seated in a primate chair with their heads firmly fixed via the head-post in the implant. The monkeys faced a LED screen (60 Hz refresh rate) 60 cm away which spanned ± 30° vertical and ± 45° horizontal of the central visual field. The position of the left eye was sampled at 500 Hz by an EYELINK1000 infrared eye tracker (SR Research). Behavioral tasks were under the control of Monkeylogic software (http://www.monkeylogic.net/). All data analyses were performed offline on MATLAB (version R2014a, Mathworks Inc).

### General procedures

Both single-neuron recording and subthreshold micro-stimulation were performed in the intermediate and deeper layer of the SC (between 1.0 and 3.0 mm below, and tangential to, the surface of the SC).

Tungsten microelectrodes (FHC Inc.) were lowered with a Microdrive (NAN Instruments) through 23 gauges, 42 mm long stainless steel guide tubes (with 10 mm spacer) attached to a Crist grid (Crist Instruments Co., Inc.). Single-cell discharges were sampled at 22 kHz using the AlphaLab SnR System and subsequently offline sorted using Spike 2 (CED, Cambridge Electronic Design Limited).

### Behavioral tasks

Monkeys performed two behavioral tasks. The delayed saccade task was used to characterize neuronal properties and map their response fields (RF). The saccade foraging task was used to correlate neuronal activities to value and choice. Separately, we applied stimulation during the saccade foraging task to determine how perturbing SC activity would alter foraging choices.

#### Delayed saccade task and response field identification

The delayed saccade task was performed before the saccade foraging task in both neuronal recording and micro-stimulation blocks^47^. Trials started when a fixation point appeared at the center of the screen. After monkeys acquired and fixated the fixation point for 1000 ms, a single target point appeared in the periphery for a random delay period (600-800 ms). At the end of the delay period, the fixation point disappeared which cued the monkey to initiate a saccade towards the peripheral target. Monkeys received a liquid reward if they initiated a saccade within 1000 ms and maintained fixation within 3 degrees of the target and held it for 300 ms. Both the fixation point and target stimulus shared the same luminance and size (4.20 cd/ m^2^ luminance, 0.5° visual radius) as targets in the saccade foraging task.

The center of an isolated single neuron’s (recording experiments) or multi-units’ (stimulation experiments) RF was defined as the location relative to central fixation that was associated with the most vigorous activity following target appearance and during target-directed saccade in the delayed saccade task. For stimulation experiments, we also used supra-threshold stimulation to verify that the stimulation-induced vector and recorded RF vector displayed close correspondence. Our experiments focused on SCi sites with smaller saccade vectors ranging from 3-13° to allow an array of targets to fit on the screen during the saccade foraging task.

#### Saccade foraging task

In the saccade foraging task (Fig.1), each trial began with the display of a grid array of circular stimuli on the screen in front of the subject. The grid was rotated, scaled and the number of targets adjusted (3×3, 4×3, or 4×4) such that when fixating a target, a nearby target would be located in the center of the neuron’s RF (except, of course, when fixating the outermost contralateral column). All visual targets were identical in terms of luminance and size (4.20 cd/ m^2^ luminance, 0.5° visual radius), but could either be red, green or blue. During each trial, target colors were represented in equal proportions, but their locations on the grid were randomly shuffled between trials. Throughout a block of trials, visual targets that shared the same color were associated with the same value as defined by reward magnitude divided by fixation time (value = reward magnitude / fixation time; see Supplementary Table 1). That is, monkeys harvested a specified volume of liquid reward associated with a colored target by fixating it for a particular duration. Once harvested, the target’s color turned into an equiluminant grey and could not be harvested again during that trial. Prior to each block, both reward magnitudes and fixation times were selected randomly with replacement while excluding identical combinations. Monkeys were free to harvest targets in any order they chose. However, we adjusted trial durations such that there was not enough time to harvest all targets. On average, 79 ± 6% of the targets were harvested.

In contrast to the recording experiments, only two value associations were used in the stimulation saccade foraging task: the similar-value condition and the different-value condition. Under the similar-value condition, reward magnitude (μL) / fixation time (s) associations were 30:1, 45:1.5, and 60:2, such that the three colors shared the same objective value of 30 μL/s. Under the different-value condition, reward magnitude (μL) / fixation time (s) associations were 20:2, 40:1.5, and 60:1, such that the values associated with the 3 colors were 10, 27, and 60 μL/s, respectively. The results from both value conditions were consistent, therefore we analyzed them together.

### Behavioral analysis

#### Rank value

The rank value quantified the relative preference for each color using the order in which each target was chosen (e.g., Fig. 2b and c). Importantly, this measure does not provide an exact measure of the subjective value of each color but it does provide an ordinal rank value for each color. This measure is derived from the nonparametric Kruskal-Wallis test^48^. The first chosen target in a given trial was given a score of N, where N is the total number of targets initially presented in the grid. The next chosen target was given a score of N-1 and scores decreased by 1 each time until the last choice:

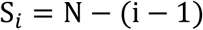

Where S_*i*_ is the score of the ith chosen target.

All targets that were not selected by the end of the trial were given equal scores that was the mean of the scores that they jointly occupy:

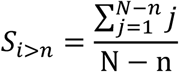

Where n is the number of targets harvested, S_*i*>*n*_ is the score of each target that was not selected by the end of the trial and j is each of the remaining scores.

The rank value (RV) is the summed score of that color (∑ *S*_*C*_) normalized by dividing by the summed score of all colors. Because the number of targets cannot be equally divided amongst the 3 colors in the 16-target conditions, we adjusted the equation by dividing the summed score of each color (∑ *S*_*C*_) by the number of targets of that color (N_C_).

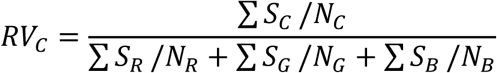

Where C is one of the colors R, G, or B, and S_*C*_ represents the score of each target that features the color C.

Sliding average of *RV*_*C*_ over five trials was used to reduce the trial-by-trial variability. The *value ranking* was ordered by the average *RV*_*C*_ in a block.

#### Time to behavioral acquisition

To quantify when the subjects learned the color-value associations, we did pairwise comparison between the rank values across the three colors within a moving 5-trial window with 1-trial steps (Kolmogorov-Smirnov Test). Time to behavioral acquisition was the trial when the RV_C_ was significantly separated in order of value ranking of the block, and remained so for at least 5 consecutive trials (e.g., Fig. 2b-e).

#### Efficiency

We defined the efficiency, *E*_*i*_ of each choice (e.g., Fig. 2d and e) as the value associated with that choice, *V*_*i*_, divided by the highest available value present at the time that the choice was made, *V*_*Hi*_:

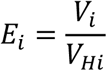

The foraging efficiency associated with each trial was defined as the sum of the efficiencies of all the choices made in that trial, divided by the number of chosen targets or, in other words, the average efficiency of the trial:

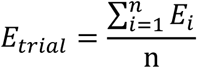

Where *E*_*trial*_ denotes the foraging efficiency of the trial, *E*_*i*_ denotes the efficiency of the ith choice, and n denotes the number of choices made in the trial.

To examine if subjects’ efficiency was significantly higher than random choices, for each block the chance efficiency was computed by simulating a random selection process over 5000 iterations. Only blocks with significantly higher efficiency than chance efficiency were included in subsequent analyses. Trials before behavioral acquisition were not included in the population efficiency calculation.

#### Behavioral criteria

Data sets had to satisfy three behavioral criteria to be included in further analyses.

1. Only trials after behavioral acquisition were analyzed.
2. Only trials in which both the order of single-trial RV_C_ and sliding RV_C_ were consistent with the overall value ranking of the block were included in subsequent analyses. However, entire blocks were removed from analysis if there were more than 25% inconsistent trials within a block (i.e., unstable blocks). One exception to this rule occurred during some similar-value, stimulation blocks when preferences occasionally switched for extended periods (at least 10 trials). Analyses followed these extended preferences rather than average value preferences across the block.
3. Lastly, total useable trials within a block must exceed 50 for recording blocks and 100 for stimulation blocks.

Based on these criteria, 70 of the 96 blocks in recording experiments met the behavioral criteria and 72 of the 78 stimulation experiments met the behavioral criteria and were used in subsequent analyses.

### Neuronal analysis

We recorded 96 neurons in the intermediate and deeper layer of SC. For the saccade foraging task, the peri-stimulus time histograms (bin width 30 ms, 15 ms steps) were aligned on the beginning of fixation, reward delivery and saccade onset. Each neuron’s activity was normalized by the average neuronal responses during the last 300 ms of fixation period (300 ms before reward delivery) when a valued target was in the RF. We divided neuronal activities by the normalization factor to get the normalized response for each neuron. If the normalization factor was less than 5 spikes/s, the neuron was defined as unmodulated by the task and would be excluded from remaining analyses. Based on the behavioral exclusion criteria described above in addition to the normalization criteria, a total of 53 single neurons (monkey D, 24; monkey R, 29) were included in our recording analyses.

#### Multiple linear regression analysis

We generated a general multiple linear regression model to assess the contribution of value and choice direction on the neuronal responses. Only the first and second value rankings were used in the menu of 3-values remaining condition because monkeys rarely chose the 3^rd^ value ranking targets. When only 3rd value ranking targets remained, all the targets were associated with the same value, therefore only the factor of choice direction was used.

We used the following regression model to fit the neuronal responses:

FR(t)=b0(t)+b1(t)*Value+b2(t)*Choice;

Where FR represents the firing rate, Value is the value ranking (either 1 or 0 representing the higher or lower value ranking, respectively) and Choice is the direction of the next saccade (either 1 or 0 depending on whether the next saccade was directed into or out of the response field, respectively). As these factors are in the same range (0 to 1), it allows us to compare directly the values of the fit coefficients (b1 and b2) and determine their impact on FR.

Statistical significance was determined with a t test with a false discovery rate (FDR) correction^49^.

#### Receiver-operating characteristic (ROC) analysis

ROC analysis was used at each time point to investigate how the decision process evolved for each value ranking^50, 51^. ROC curves were derived from the distributions of choice-in activities and choice-out activities of the same value ranking within a given menu (Fig. 4c).

### Electrical stimulation experiments

The parameters for electrical stimulation were determined using the delayed saccade task. Clear delay-period multi-unit activity must be detected before stimulation to ensure stimulation sites were comparable to recording sites. To find the stimulation threshold, 0.25 ms biphasic currents (10 ms duration, 300 Hz) were applied during the delay period and systematically reduced from 30 μA until only hypometric saccades could be triggered. We set this intensity as the threshold intensity (R, average 13.6 μA; D, average 19.3 μA). During the stimulation saccade foraging task, we increased the stimulation duration to 300 ms and decreased the frequency to 150 Hz.

Subthreshold stimulation was randomly applied on half of the fixations on 2/3 of the trials. The remaining 1/3 trials were control trials. Stimulation trials and control trials were randomly interleaved. Stimulation was applied in the last 300 ms of the fixation period before reward delivery. The stimulation bias was the proportion of saccades directed towards the stimulation site for stimulation trials minus non-stimulation trials in every block.

## Acknowledgment

This work was supported by grants from the CAS Hundreds of Talents Program. We thank M. Yang, X. Wang, G. Yu, H. Liao, S. Xie for their help in all phases of the study and Z. Zhang, C. Duan, T. Yang for constructive feedback regarding the manuscript.

## Author Contributions

B.Z., J.Y.Y.K. and M.C.D. conceived the project and designed the experiments. B.Z. performed the experiments and analyzed the data. B.Z. drafted manuscript, B.Z., J.Y.Y.K. and M.C.D. edited and revised manuscript.

---

**Supplementary Figure 1.**
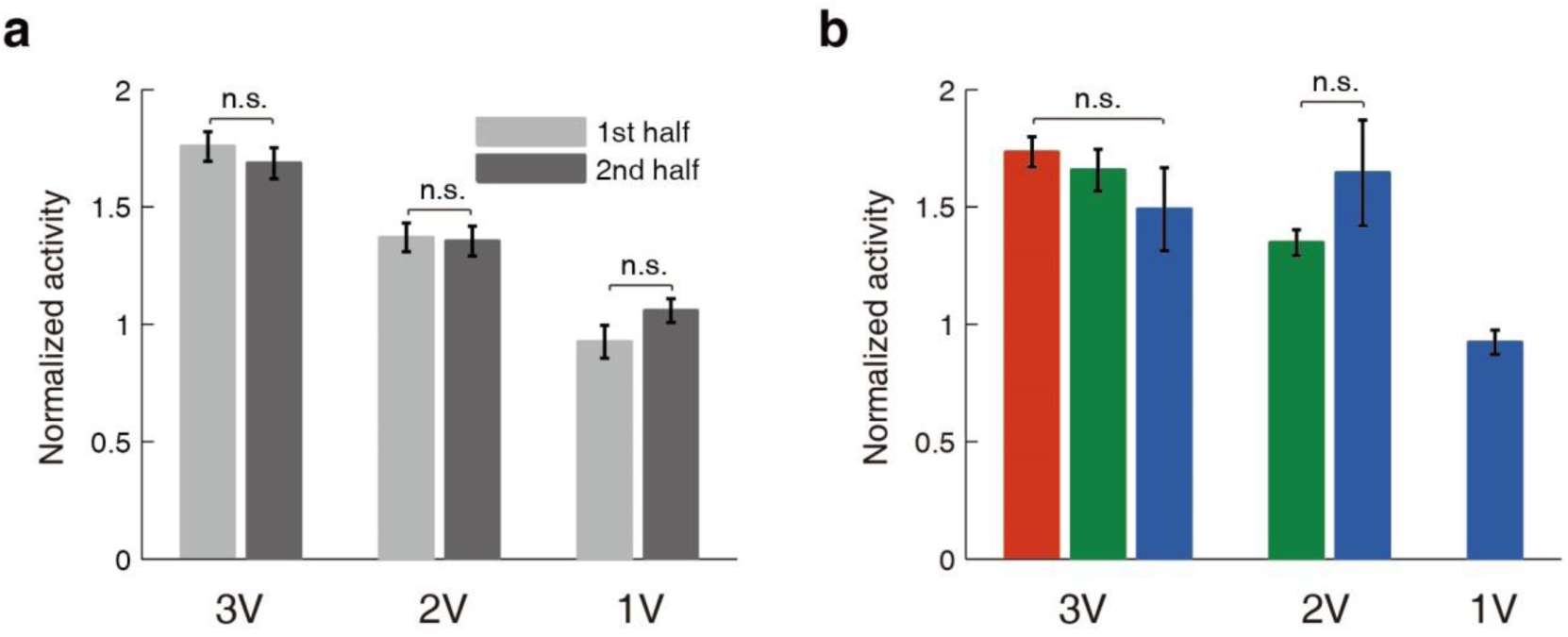
Factors that had no effect on selection level. (a) For each menu, the selection level did not change whether a value item was chosen early (light gray – 1^st^ half of value items) or late (dark gray – 2^nd^ half of value items) in the trial sequence. (Paired-t tests) (b) The value of the fixated targets had no effect on the selection level within a given menu. Data is presented as a function of the value of the fixated target when there were three values (3V), two values (2V) or one value (1V) targets remaining in the array. Error bars represent ± SEM. (One-way ANOVA test and Paired-t test)

**Supplementary Figure 2.**
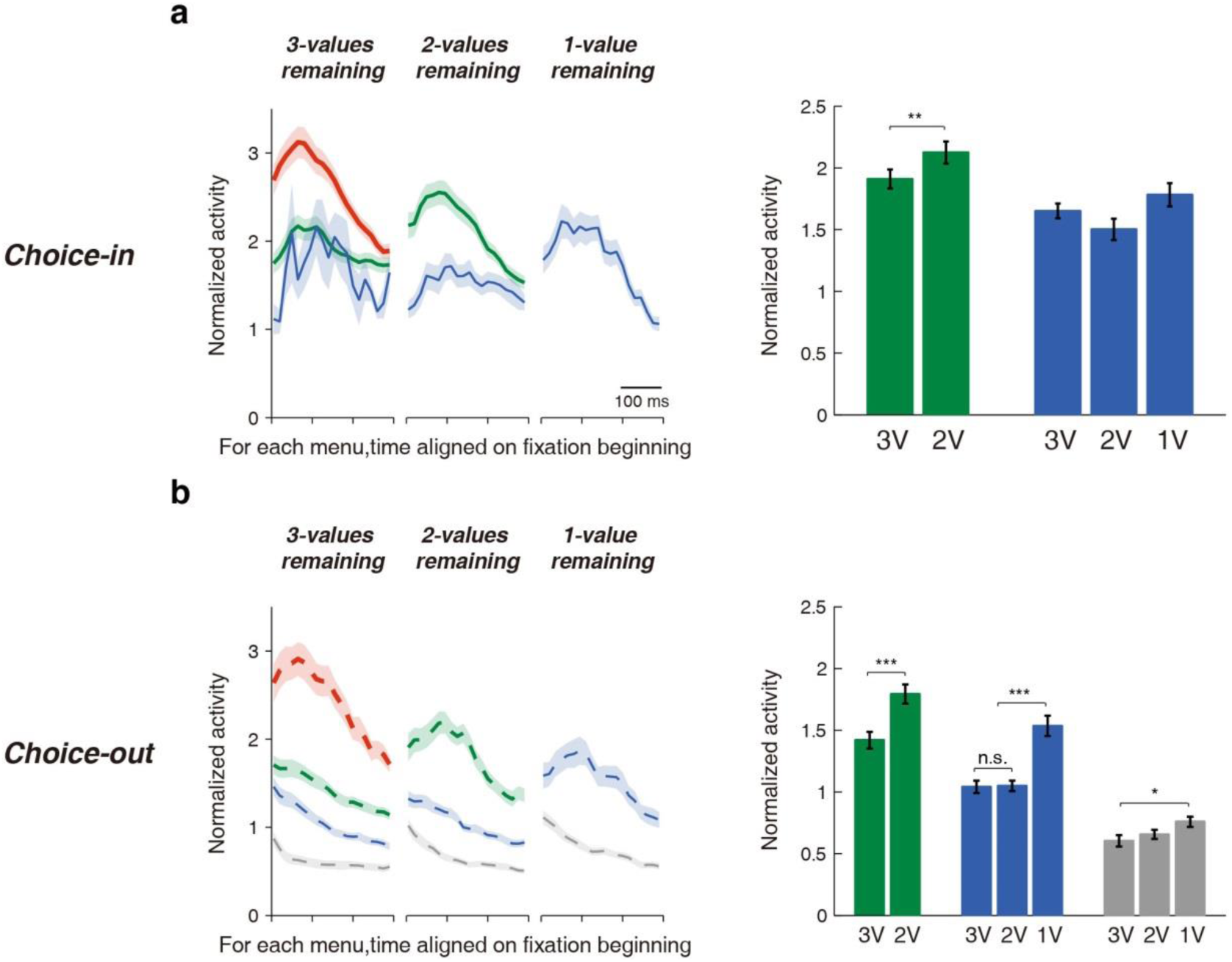
Menu-dependent SC early fixation period neuronal activities. Normalized neuronal activity in the first 300ms of the fixation time from choice-in (a) and choice-out conditions (b). Same format as Fig. 5. Data is presented for each value ranking when there were three values (3V), two values (2V) or one value (1V) targets remaining in the array. Error bars represent ± SEM. (Paired-t tests and one-way ANOVA tests, *post hoc* tests were done with Bonferroni correction)

**Supplementary Figure 3.**
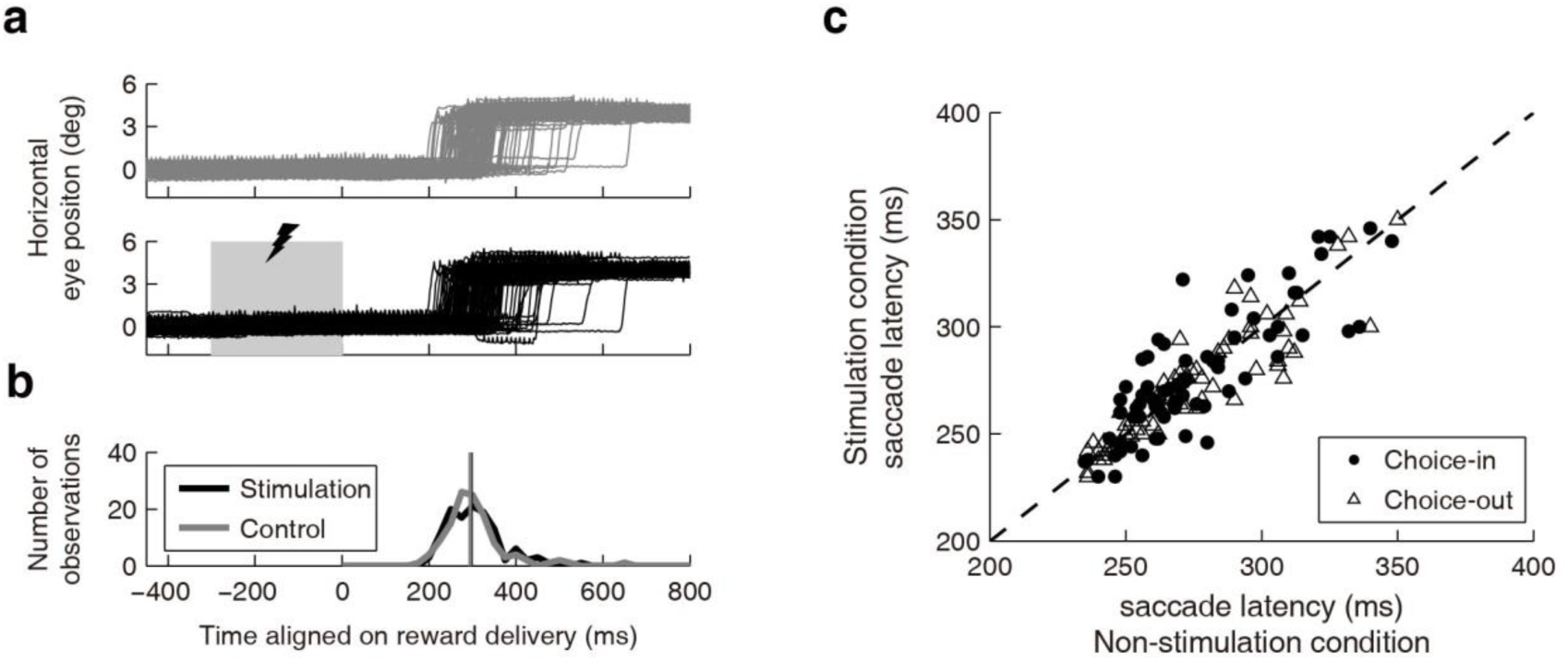
Micro-stimulation was sub-threshold for directly triggering saccades. (a) Horizontal eye position for saccades directed towards the stimulation site for an example block of trials. Gray traces denote non-stimulation, control conditions and black traces denote stimulation conditions. The shaded gray area and associated lightning bolt indicate the period during which micro-stimulation was applied. The time is aligned on reward delivery. (b) The corresponding saccade latency distributions for the example block. Vertical solid lines indicate the median latency of the distributions. (c) Median saccade latency for each stimulation site during stimulation compared with non-stimulation condition. Choices toward the stimulation site are denoted by circles and choices out of the stimulation sites are denoted by triangles.

**Supplementary Table 1.** Possible target values during neuronal recordings

| Value ($\mu\text{L/s}$) | | Fixation time (s) | | | | |
| --- | --- | --- | --- | --- | --- | --- |
|  |  | 0.5 | 1 | 1.5 | 2 | 2.5 |
| Reward magnitude ($\mu\text{L}$) | 20 | 40 | 20 | 13 | 10 | 8 |
|  | 40 | 80 | 40 | 27 | 20 | 16 |
|  | 60 | 120 | 60 | 40 | 30 | 24 |
|  | 80 | 160 | 80 | 53 | 40 | 32 |
|  | 100 | 200 | 100 | 67 | 50 | 40 |

**Supplementary Video 1** Movie for example trial of saccade foraging task (same trial as Fig. 2a). The white dot represents foveal eye position. The gray circle denotes the location and approximate size of the SC neuronal response field that is represented in retinotopic coordinates and consequentially moves in concert with the eye. The auditory beeps represent the timing and duration of liquid reward delivery. The timing of neuronal spikes is represented by the white raster and associated auditory tick. The duration of fixation and mean firing rate associated with different colored targets in the neuron’s response field are denoted by the vertical and horizontal colored bars at the bottom, respectively.

